# Proteogenomic analysis of the differential stability of cardiac protein isoforms

**DOI:** 10.64898/2026.07.23.740363

**Authors:** Matthew Juber, Shaonil Binti, Boomathi Pandi, Edward Lau, Maggie P.Y. Lam

## Abstract

Alternative splicing is an important regulatory layer in gene expression, but knowledge on the isoform protein molecules continue to lag their canonical counterparts. An open question is whether alternative protein isoforms feature different half-life than the canonical counterpart, which could indicate differential usage and functional diversification. Here we combined a proteogenomics approach with heavy water-based protein turnover analysis to survey 24 pairs of canonical-alternative protein isoforms in the mouse heart. The results provide a reference on their numerical half-life and also reveal widespread differences in isoform stability.

## Introduction

Alternative splicing and transcript usage is broadly implicated in heart diseases and transcript isoform heterogeneity is widespread in the cardiac system.^2^ For instance, recent work suggests that reverse cardiac remodeling after ventricular assist devices is primarily associated with splicing rather than gene expression changes including an isoform switching of CAMK2D, reinforcing the notion that fine-grained regulations exist at the isoform levels even for assumed maladaptive genes.^3^ However, although transcriptomics data can identify which isoforms are produced, it cannot predict the molecular function and longevity of protein isoforms. Because non-canonical proteins often harbor sequence features that promote rapid degradation, the physiological significance of many splice-derived proteins remains unknown. The extent to which differential protein stability contributes to the proteoform makeup therefore remains poorly characterized.

One promising method for identifying functionally relevant splice variant proteoforms in a scalable way is to compare their turnover rates across samples. As the synthesis and degradation of proteins is an energetically expensive process, the rate of turnover is expected to be tuned to parameters including protein stability, usage, and interactions. Intriguingly, both alternative splicing differential regions as well as protein degradation driving elements (degrons) are heavily enriched within disordered regions, which has prompted efforts to evaluate how alternative splicing may affect protein half-life. Several pioneering works have attempted to compare protein isoform half-life^4–6^ though technical difficulties remain, namely:

1. It remains difficult to unambiguously resolve splice variant proteotypic proteins through unique exon/junction-specific peptides. Isoforms share most of their sequences with the canonical isoforms, hence to compare their turnover, unique peptides need to be found. Some splice variants are not in databases; whereas an overly redundant database will decimate the number of isoform unique peptides.
2. An alternative is to discover potential isoforms de novo from peptides that behave differently from the rest of the protein. This approach has been used for isoform and also post-translational modification (PTM) peptide turnover, but generally remain challenging as peptide-level turnover data can be noisy.
3. There is a paucity of stable isotope kinetics experiments performed in animal models. Most turnover studies have been done in cellular systems which have different turnover regimes than in vivo studies, and also likely different isoform expression patterns than animal tissues.
4. Most in vivo turnover studies only report single values of turnover rates or half-life per protein, without giving error estimates; and typically, statistical inferences remain underdeveloped, meaning it is unclear if any differences in protein half-life, whether between two proteins or for the same protein between two conditions/species/tissues, is statistically significant.

To bridge this gap, here we implemented an integrated analytical framework to quantify the in vivo half-lives of endogenous cardiac protein isoforms.

## Results and Discussion

To compare protein isoform half-life we leveraged D_2_O labeling mass spectrometry across six mouse strains (PXD002870)^7^ to capture proteostatic demands in their native physiological context (**Figure 1A**). To overcome the inherent difficulty of isoform-specific peptide assignment, we generated an RNA-seq guided database using a software we previously developed, JCAST^8^, to re-search large data sets of D_2_O labeled experiments. JCAST translates sequences from mRNA data, and so can identify potential tissue-specific isoforms that are not found in protein sequence databases. Even for well annotated organisms, JCAST may be useful for filtering out absent transcript isoforms that are unlikely to present as proteins, which helps find isoform-unique peptides.

**Figure 1.**
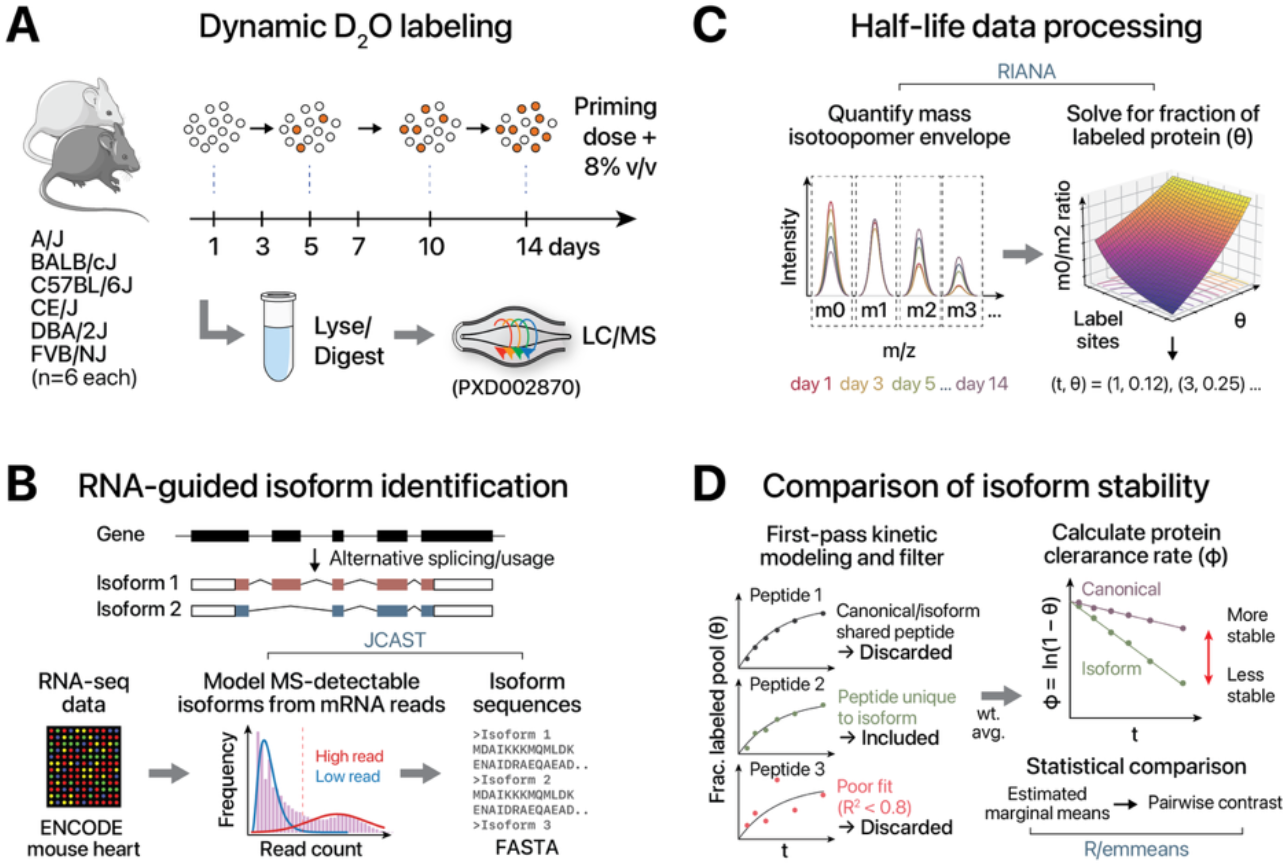
Analytical workflow to process D_2_O labeling data and calculate the stability of canonical proteins and their isoforms.

Next, we integrated the proteogenomics-derived custom databases with a custom protein parsimony procedure that is isoform aware and sits in between gene-level parsimony and unique peptides in selectivity. Briefly, peptides that map to two different gene names are discarded from further consideration, as it cannot be ascertained whether the isotope incorporation time-course data comes from which gene. Peptides that map to two different protein entries of the same gene name may be retained. If the peptide maps to both canonical and an alternative isoform, we assume that the peptide emits from the canonical isoform, unless the alternative isoform is supported by an additional unique peptide, in which case the peptide is discarded. This is based on the logic that most genes have one dominant isoform, and that this abundance isoform is typically the canonical form of the protein. Thus, while most alternative isoforms may only differ from the canonical isoform for say one exon and so share most of its sequence, the shared sequence does not provide independent evidence of the alternative isoform’s evidence, and is grouped under the canonical isoform to avoid removing most peptides from consideration (**Figure 1B**).

To analyze the isotope incorporation data, we use an analysis routine we recently described to improve the depth and estimation of kinetics parameters from D_2_O mass spectrometry data^9^ (**Figure 1C**). To address the need for rigor in turnover studies, we recently applied a linearized first-order kinetic model to estimate protein half-life and infer inter-group significance. A first-order kinetics model can be linearized by taking a log transform on both sides of the equation to provide an easy mean to estimate the turnover rate coefficient *k* from linear models (Methods). While this method is well established in the literature^10,11^, it is uncommonly applied in proteome-wide turnover studies which often compare groups using peptide observations as pseudo-replicates. Here, we extend the linear model method to half-life comparison within an isoform group, by performing statistical testing via estimates of marginal means and trend contrasts then Tukey’s adjustment (within-group) and Bonferroni correction (across-groups) (**Figure 1D**). This method allows rigorous comparison of protein half-life across groups using biological replicates rather than pseudoreplicate measurements from multiple peptide observations within each protein.

Using this framework, we achieved unambiguous assignment and resolved the half-lives for 24 canonical-isoform pairs. Measured canonical protein half-lives span 1.1–17.7 (median 4.3) days; vs. 1.5–9.1 (median 3.9) days in alternative isoforms. Consistent measurements are observed across mouse strains, indicating protein half-life is conserved across genetic backgrounds. Our analysis reveals that protein stability is highly regulated at the isoform level. Whereas some pairs share near-identical stability (e.g., the full-length obscurin and truncated obsc-80 isoforms) (**Figure 2A**), others have different stability. For instance, spectrin alpha chain, non-erythrocytic 1(SPTAN1)’s isoform-2 has significantly different half-life than the canonical isoform (P: 2.3e–6) (**Figure 2B**). Myosin light chain 1/3 isoform MLC1 (A1) have half-life of 7.7 days vs. isoform MLC3 (A2)’s 1.5 days (P: 2e–9) (**Figure 2C**). These two well-described isoforms differ by the N-terminus from alternative translation start sites due to the first two exons. The A1/A2 forms are known to differ in actin-myosin interaction and contraction kinetics; hence the results are consistent with half-life reflecting usage and functional differences. Sorbin and SHR domain-containing protein 2 (SORBS2) presents an example where a group of 3 isoforms have different half-life to each other (P: 5.0e–9 between isoform-5 and canonical; P: 1.3e–7 between isoform-2 and canonical) (**Figure 2D**).

**Figure 2.**
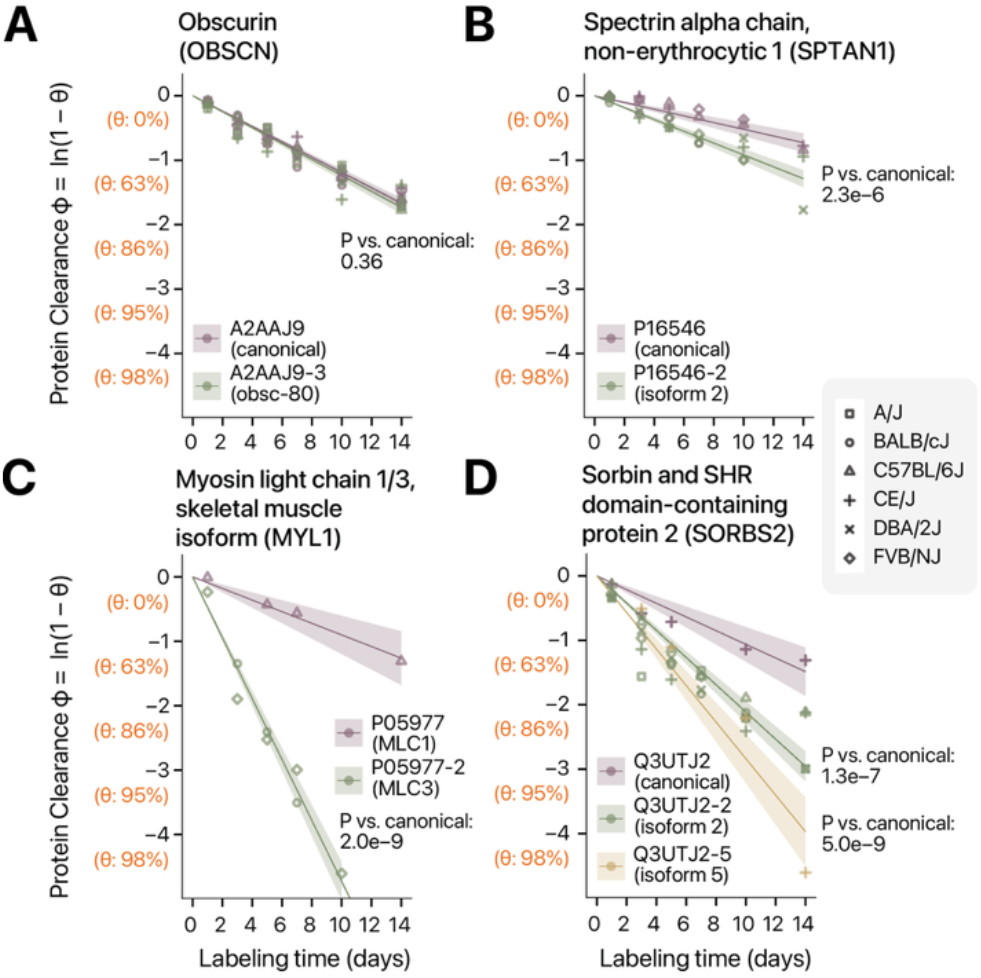
Protein stability curves of canonical/isoform protein pairs with similar (**A.**OBSCN) and significantly different (**B**. SPTAN1, **C**. MYL1, **D**. SORBS2) stability. X-axis: D_2_O labeling time (days); Y-axis: protein clearance ϕ.

On a proteome level, a striking 63% (15/24) of the queried isoforms exhibit significantly different turnover rates compared to their canonical counterparts (Bonferroni P<0.05) (**Table 1**). This differential stability is distributed across key functional categories, including the sarcomere, mitochondrial metabolism, and calcium signaling. Notably, in 60% of these cases, the canonical isoform was more stable than its alternative counterpart. The preponderance of unstable alternative isoforms suggests a possible proteome architecture that primes transient alternative isoforms for rapid proteomic shifts, while relying on stable canonical isoforms to maintain structural integrity. On the other hand, while most genes are thought to have one prominent, highly expressed isoforms, the results suggest that stable alternative isoforms are readily identifiable in the proteome. Stable protein isoforms are also quantified from JCAST that are not found in the de facto standard protein database UniProt Swiss-Prot, mapping to Ensembl Eno1-205, Calu-202, Actn1-202 transcripts.

**Table 1.**
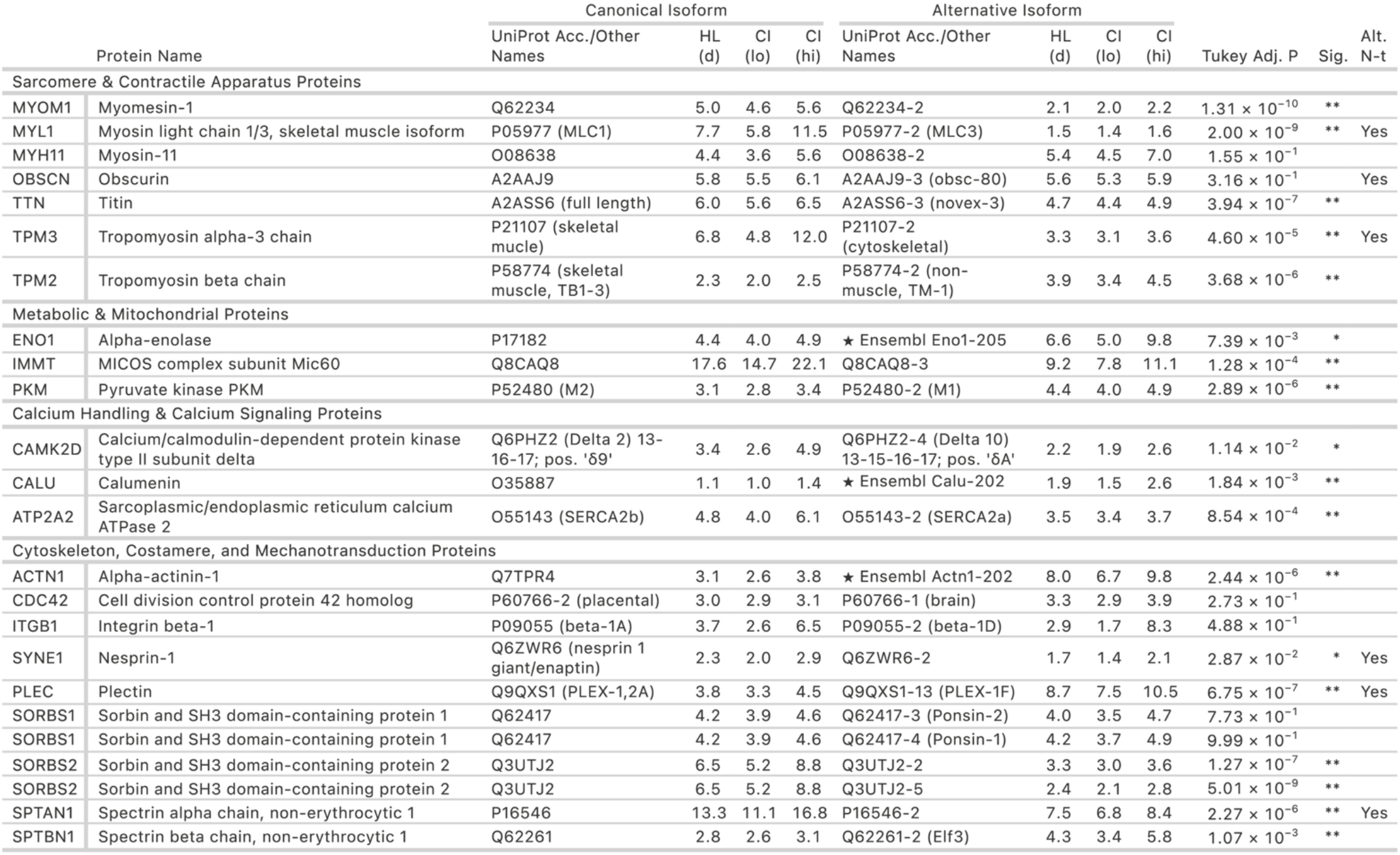
Table of 24 canonical-alternative isoform pairs with unambiguous peptide assignment and confident protein stability calculations. ★: RNA-seq translated sequences, corresponding to Ensembl transcripts; *: P < 0.05 with Tukey’s HSD; **: P < Bonferroni corrected P threshold across proteins. Alt. N-t: isoform contains alternative N-terminus sequence.

Lastly, we examined instances where stability shifts are associated with specific sequence motifs elements by querying documented degrons in Degronopedia^12^. Interestingly, 23 out of the 24 canonical-alternative isoform pairs feature at least one candidate degron sequence motifs that is unique to either the canoncial or the alternative isoform sequence alone, suggesting there is broad opportunity for differential degradation regulations (**Table 2**). In multiple instances, the annotated primary degron motif sequences (e.g., SORBS2 isoform -2 motif SPP 355–357) lie adjacent to ubiquitinated lysine sites and are located within long intrinsically disordered region (IDR) boundaries, making them likely candidate degradation regulator in the tripartite degron model. In other instances, the isoform-unique primary degron motif sequences lie adjacent to the accessible protein N-terminus and may be substarate of the N-degron pathway (formerly N-end rule). For example, the TPM3-2 isoform possesses an N-terminal “MAG” sequence absent in the canonical form, potentially subjecting it to the Ac/N-degron pathway via Ccr4-Not4 (**Figure 3A**).

**Table 2.**
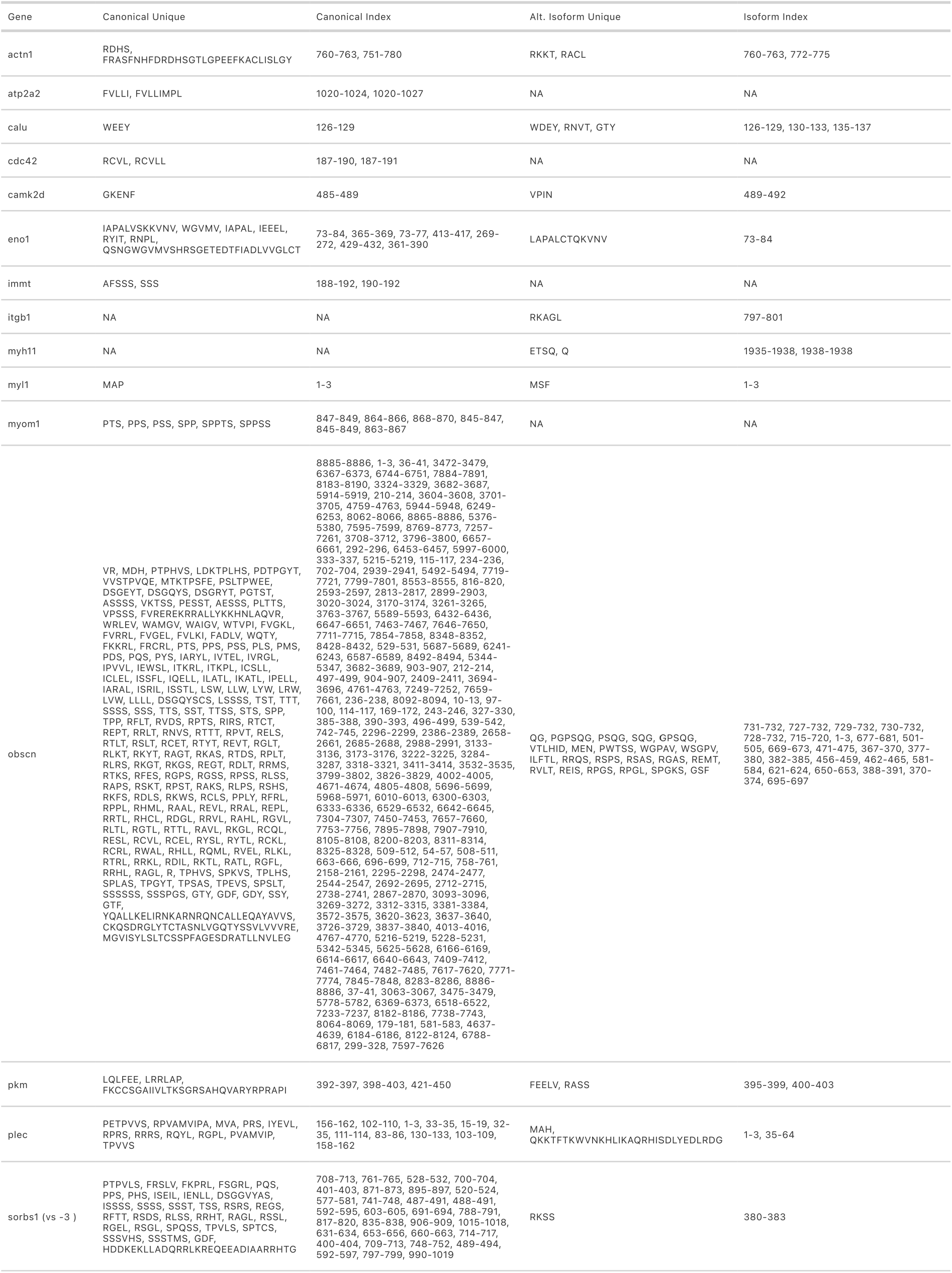
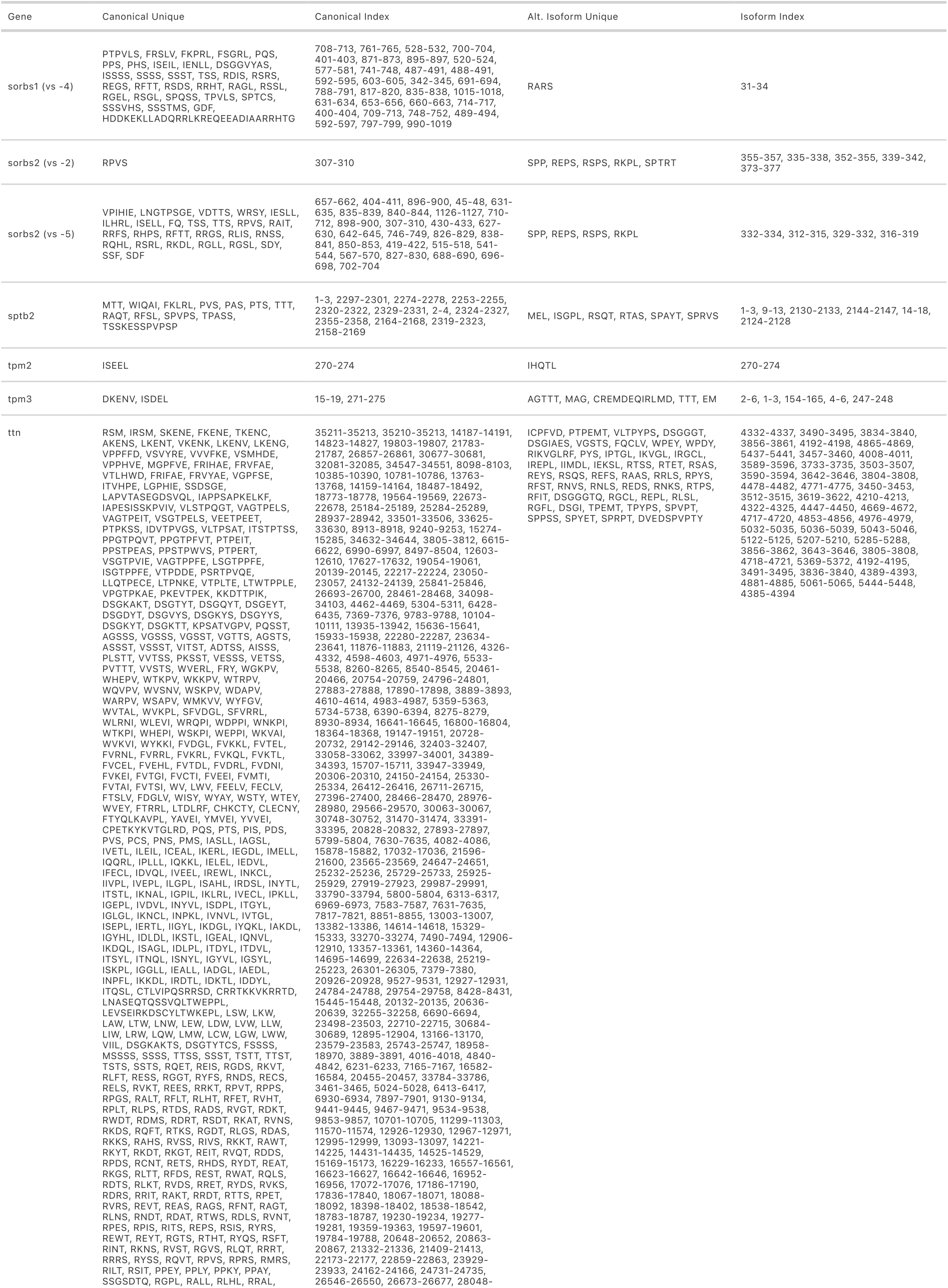
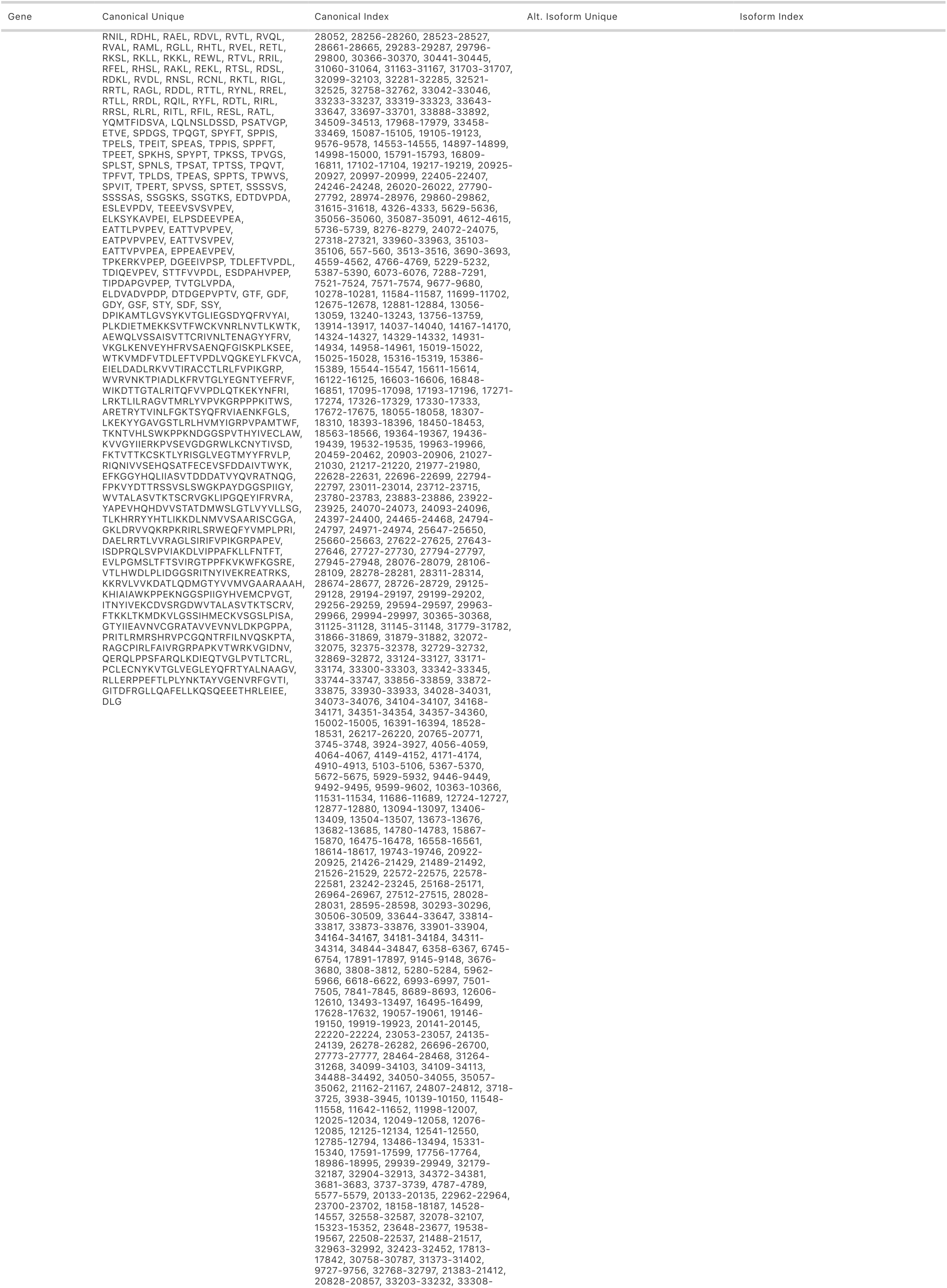
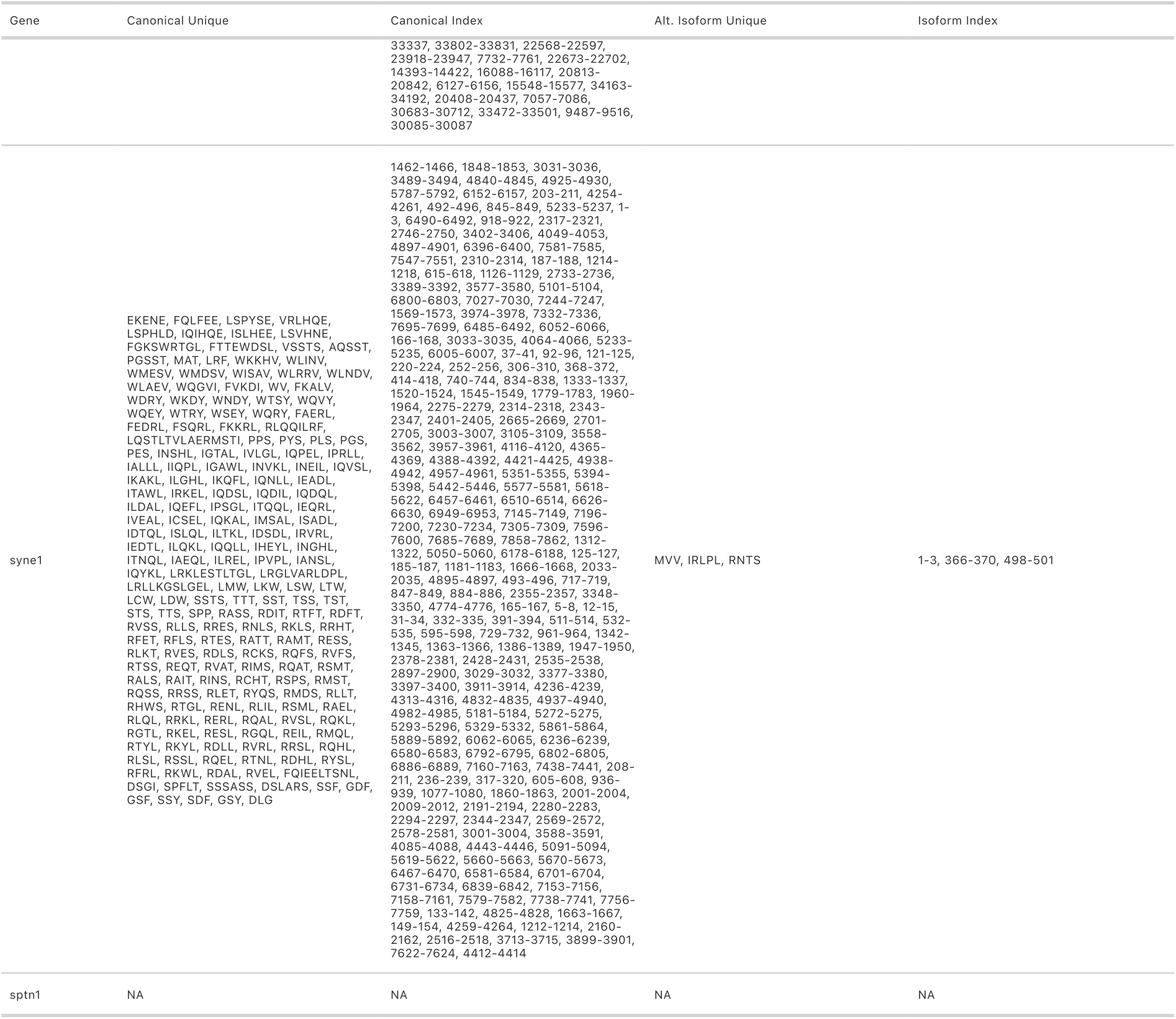
Table of candidate degron motif differences between canonical and alternative isoform pairs. For each gene, found instances of unique Degronopedia annotated candidate degron motifs (in cases where multiple identical motifs exist, only one representative instance is kept), in each of the canonical and the alternative isoforms are shown.

**Figure 3.**
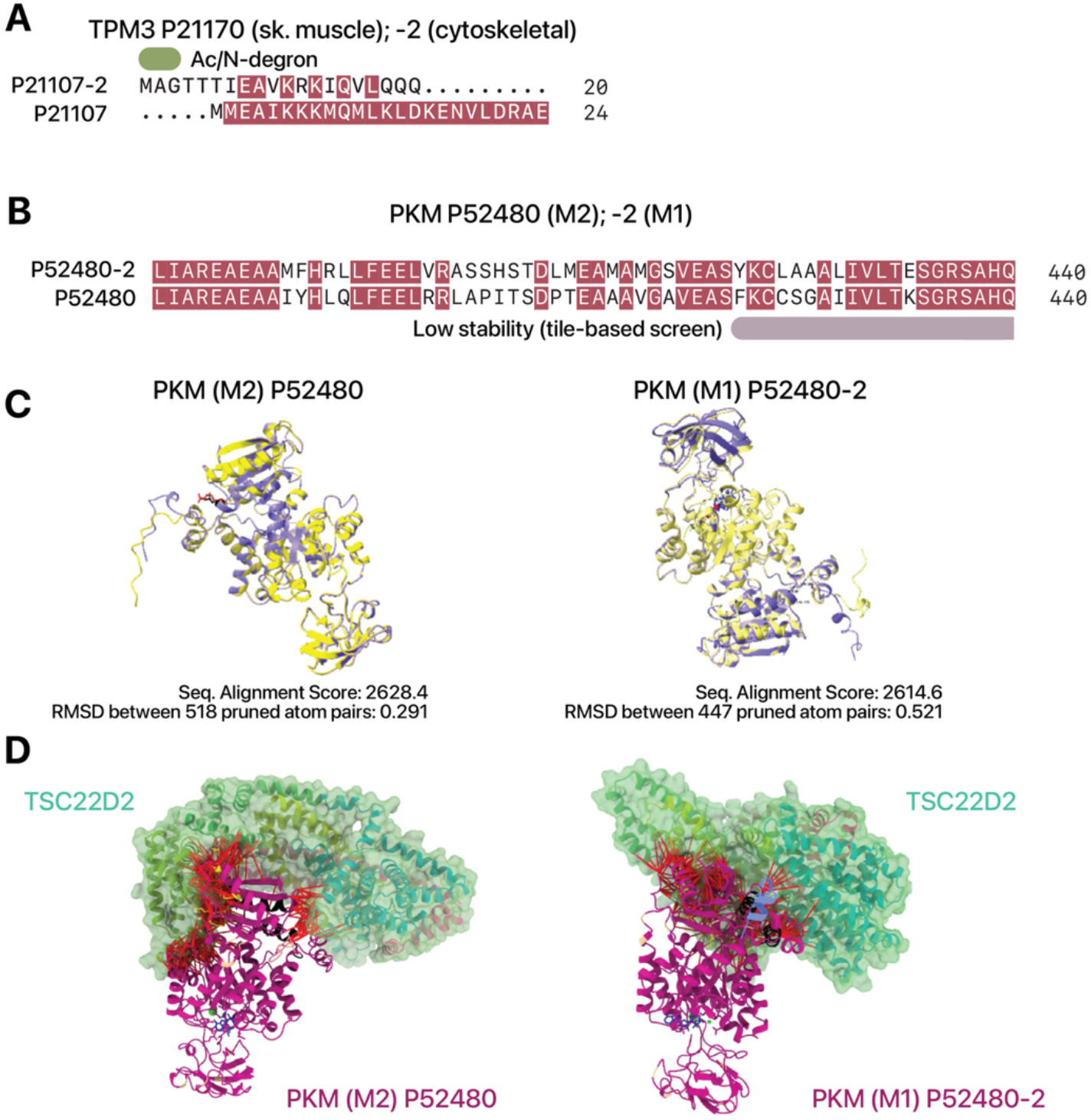
**A–B:** Aligned sequences and degrons for **A**. TPM3 and **B**. PKM isoforms. **C**. AlphaFold3-predicted structures of PKM M2 and M1. **D**. Predicted interactions with TSC22D2. Red lines show interactions between chains, with the two isoforms differing in predicted interactions with TSC22D2 at residues 432–437, 482–489, and 516–521.

In another example, the PKM M2 isoform differs from M1 via a mutually exclusive exon containing a stretch of residues (421-450). This sequence is annotated to be degradation relevant on Degronopedia as it is reported to confer low in-vitro stability in a large protein-tiling screen^13^, suggesting it may act as a standalone degradation/stability regulator element (**Figure 3B**). In parallel, while further experimental validation is needed to confirm finings, we also explored AlphaFold3 (ref ^14^) predicted structures of PKM M2 and M1 with the known PKM interactor TSC22D2, which show different interactions including from allosteric interactions beyond the isoform-variable sequences (**Figure 3C–D**). Taken together, these analyses suggest distinct possibilities for isoform-specific sequences and structures may contribute to differential stability.

To summarize, this report represents to our knowledge the first systematic survey of differential protein stability among splice isoforms in an animal model. We provide a straightforward literature record of the endogenous half-life of 24 pairs of proteins and isoforms known to be expressed in the mouse heart. Despite the depth of this study being limited by the source data, our findings demonstrate that isoform switching carries proteostatic implications, where the regulation of protein lifetime occurs in parallel to protein sequence usage. This study also provides a methodological framework for comparing proteome wide alternative isoform-specific stability in other models.

## Methods

Mass spectrometry data were retrieved from ProteomeXchange accession PXD002870^7^, which contained proteomics data measured from the cardiac tissues of six strains of mice labeled with up to ∼5% body water excess deuterium and collected at 1, 3, 5, 7, 10, and 14 days post labeling. The data were then researched with MSFragger (v4.3) within FragPipe (v23.1) ^15^ headless mode against UniProt Swiss-Prot protein sequences appended with additional isoform sequences ^16^. Briefly, the mouse heart isoform specific database was generated using JCAST v0.34 using short-read C57BL6/J mouse heart RNA-seq data as described ^17^. The translated isoforms were then harmonized against UniProt Swiss-Prot mouse canonical and isoform sequences retrieved on 2025-09-22. Precursor and fragment mass tolerances were set to ±20ppm. Digestion parameters were set to strict trpysin (K/R), with up to 2 missed cleavages allowed. Modification parameters were as follows: fixed Cys carbamidomethylation (+57.02146 Da), variable: Met oxidation (+15.9949 Da) and protein N-terminal acetylation (+42.0106 Da), ≤3 per peptide. Isotope error was set to 0/1/2/3. Peptide-spectrum matches were rescored with MSBooster (v1.3.17) before Percolator (v3.07.1) filtered peptides and proteins to <1% FDR^18,19^.

To quantify D_2_O label incorporation, the individual mass isotopomer relative abundances within a peptide envelop are integrated using RIANA v.0.8.2 as previously described ^9^. For each peptide at each time point, the fraction of newly synthesized proteins (*θ*) are numerically solved by minimizing the RMSE of isotopomer distributions between the experimental and theoretical mixture envelopes as described ^9^. A first-pass curve-fitting is then performed where the time-series data of *θ* is fitted to a first-order kinetic model to find the best-fit k for each peptide-charge combination, where for each peptide-charge combination

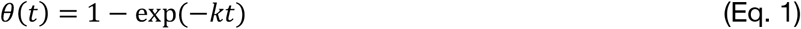

with fixed endpoints *θ*(0) = 0 and a plateau of 1. Peptides that map uniquely to a protein and that pass a goodness-of-fit R^2^ ≥ 0.8 filter of the first-pass fitting are then rolled up into proteins. We then calculate a protein clearance parameter, *ϕ* via log transform of the first-order kinetics equation:

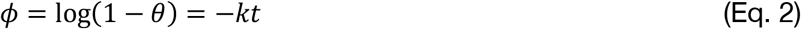

Replicate mass spectrometry observations across injections and spectra are collapsed such that the linear model only considers the variability of individual animals. As *ϕ* is linear to *t* with a slope of −*k*, a linear model and ordinary least square method can be used to estimate *k*. Moreover, by labeling all isoforms within a gene name, a joint model across isoform groups (*ϕ* ∼ 0 + day + day:isoform) with fixed intercept and shared residual variance can be used to simultaneously estimate isoform-specific slopes, confidence intervals, and contrast statistics using estimated marginal means, using EMMEANS/EMTRENDS in R. Multi-group pairwise comparison with Tukey’s HSD adjustment on P values. Family wise error rates are then controlled using the Bonferroni correction. Comparison results with Tukey’s HSD P < 0.05 are considered suggestive/trending, and P values below Bonferroni corrected threshold (0.05/# isoform groups) are considered significant. For degron analysis, protein sequences are queried using Degronopedia ^12^ to retrieve primary degron motif sequences, ubiquitination lysine positions, average sequence disorder, and distance to long intrinsically disordered regions.

## Acknowledgments

This work was supported in part by NIH grants R35GM146815 (E.L.) and NIH grants R01HL141278, and R01GM144456 (M.P.Y.L.) This work utilized the Alpine high performance computing resource at the University of Colorado Boulder. Alpine is jointly funded by the University of Colorado Boulder, the University of Colorado Anschutz, and Colorado State University and with support from NSF grants OAC-2201538 and OAC-2322260.

## Notes

### Competing Interest Statement

The authors have declared no competing interest.

